# Bias in diversity estimators and neutrality tests induced by neutral polymorphic structural variants

**DOI:** 10.64898/2026.02.26.708357

**Authors:** Sebastian E. Ramos-Onsins, Jeffrey Ross-Ibarra, Mario Caceres, Luca Ferretti

## Abstract

Estimators of genetic diversity and neutrality tests derived from the site frequency spectrum (SFS), such as Watterson’s *θ*_*W*_, nucleotide diversity *π*, Tajima’s *D*, and Fay and Wu’s *H*, are designed to be interpreted relative to a baseline defined by the standard neutral SFS. In genomic regions strongly linked to a polymorphic structural variant (SV), deviations from these baselines occur even under strict neutrality: conditioning on an SV at known frequency partitions samples into SV and non-SV haplotypes and distorts the SFS for linked neutral mutations. These deviations are well understood for genomic inversions under long-term balancing selection. However, not all SVs are under strong selection, and the evolution of some SVs may be better approximated as neutral. Here we derive analytical expectations for the unfolded (and, when necessary, folded) SFS of single nucleotide polymorphisms conditional on neutral linked polymorphic SVs, including inversions, deletions, insertions, and introgressions. We use these expectations to quantify the resulting bias in standard diversity estimators and neutrality tests as a function of SV frequency and type. Finally, we discuss approaches to build corrected estimators of diversity and neutrality tests that are unbiased/centered after accounting for the presence and frequency of the SV.

## 1 Introduction

The site frequency spectrum (SFS) is the simplest fundamental summary of polymorphism data and underpins many estimators of genetic diversity and widely used neutrality tests (Achaz (2009); Braverman et al. (1995); Bustamante et al. (2001); Stephan (2010)). In practice, quantities such as Watterson’s *θ*_*W*_ (Watterson (1975)), nucleotide diversity *π*) Tajima (1983)) Tajima’s *D*)Tajima (1989)), and Fay and Wu’s *H* (Fay and Wu (2000)) are often interpreted by comparison to a neutral baseline spectrum under which their expectations are known. These baselines implicitly assume that the local SFS is well-approximated by the standard neutral model: in fact, the standardized forms of these estimators and tests are typically defined to be centered or unbiased under this model (Achaz (2009); Ferretti et al. (2010)).

Polymorphic structural variants (SVs) violate this assumption. When a genomic region is completely linked to a segregating SV, all neutral variants in that region are inherited together with the SV ancestral or derived allele, and therefore their evolutionary history is not independent from the evolutionary history of the SV (Ho et al. (2020); Wellenreuther and Bernatchez (2018)). Accounting for the SV’s frequency in the population (or its count in a sample) thus conditions the genealogies of linked neutral sites and alters the expected distribution of derived allele counts (Ferretti et al. (2018)). As a consequence, standard SFS-based estimators and tests can be systematically biased even under neutrality in regions containing SVs, and this bias may vary strongly with SV frequency and the actual type of SV (Ferretti et al. (2018); Giner-Delgado et al. (2019)).

These biases are well understood in the context of genomic inversions, where a significant amount of empirical and theoretical work has focused on polymorphic inversions maintained by balancing selection on the ancestral and inverted alleles (Ayala et al. (1971); Charlesworth (2023); Connallon and Olito (2022); Peng *et al*. (1991); Schaeffer (2008); Wellen-reuther and Bernatchez (2018). In fact, there is evidence from very different species Charlesworth (2023); Giner-Delgado et al. (2019); Joron et al. (2011)) that many inversions evolve under strong selection. However, not all SVs evolve under strong selection, and the case of neutrally evolving variants presents different modelling challenges. Furthermore, much of this progress has focused on coalescence patterns of ancestral and inverted sequences (Charlesworth (2023)). We are instead interested in the deviations of the frequency spectrum, which depend on global properties of the genealogy in the SV region. Neutral indel variants are especially challenging, since their spectrum depends on the detailed dynamics of either ancestral or derived alleles only.

In this work we derive exact analytical expectations for the conditional SFS of neutral mutations completely linked to a focal SV, considering four simple types of biallelic SVs: inversions, deletions, insertions, and introgressions. We then quantify induced biases in main diversity estimators such as *θ*_*W*_, *π* and neutrality tests such as Tajima’s *D*, and Fay and Wu’s *H*. Finally, we propose approaches to correct for these biases and build SV-aware unbiased versions of diversity estimators and centered versions of neutrality tests.

## 2 Methods

### 2.1 Sampling and conditioning under complete linkage

Consider a sample of *n* haplotypes from a neutrally evolving population. A biallelic structural variant segregates at population frequency *f* ∈ (0, 1) for the SV allele. In the sample, the SV allele has count *i* ∈ {0, …, *n*}. We can condition either on the population frequency *f* (population-conditioned) or on the sample frequency *i* (sample-conditioned); we will focus on the latter, and report results for the former in Appendix. We also assume fair sampling of both alleles of the SV; formulae for unbalanced sampling are provided in Appendix.

Throughout, we assume complete linkage: all analyzed sites are perfectly associated with the SV allele, so each sampled haplotype belongs unambiguously to the SV or reference background across the region of interest.

The conditional SFS derivations for linked sites under neutrality adopt a focal-allele conditioning strategy and corresponding finite-sample conditional spectrum objects. The analytical approach follows (Ferretti et al. (2018)) and is specialized here to focal alleles corresponding to SVs, with SV-type-specific observability rules.

### 2.2 Unfolded and folded site frequency spectra

Let *ξ*_*k*_ denote the number of neutral single-nucleotide variants (SNVs) with derived allele count *k* ∈ {1, …, *n* − 1} in the sample (the unfolded SFS). Our primary object is the *SV-conditional expected spectrum*

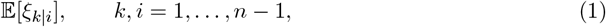

for different types of SVs.

When ancestral states are partly unknown, such as the case of insertions/introgressions with unknown origin of the inserted sequence, we use the folded spectrum *η*_*k*_ for minor allele count *k* ∈ {1, …, ⌊*n*/2⌋},

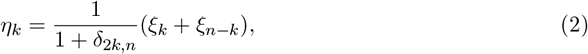

### 2.3 Bias of diversity estimators and neutrality tests

Let *T* be an SFS-based statistic. We define the *SV-induced bias* under neutrality as the deviation between the SV-conditional expectation and the baseline expectation based on the standard neutral spectrum ∝ 1*/f* (or 1*/k*):

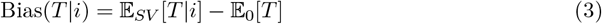

where the baseline 𝔼_0_[*T*] is the unconditional neutral expectation, i.e. the genetic variability parameter *θ* for (unbiased) estimators and 0 for (centered) neutrality tests.

### 2.4 Conditional expectations of *θ*_*W*_ and *π*

Both Watterson’s *θ*_*W*_ (Watterson (1975)) and Tajima’s *π* (Tajima (1989)) are linear functions of the SFS (Achaz (2009); Ferretti et al. (2010)). Conditional on *i*, their expectations are

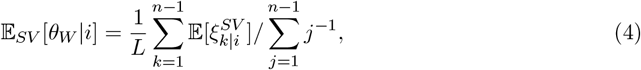

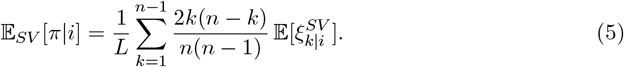

Substituting 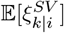 for the detailed spectra above yields explicit frequency-dependent expectations.

### 2.5 Tajima’s *D*

Tajima’s *D* (Tajima (1989)) is a standardized function of (*π* − *θ*_*W*_). When conditioning on the presence of a SV of frequency *i*, 𝔼_*SV*_ [*D*|*i*]≠ 0 in general, even under neutrality. The magnitude and sign depend on the specific conditional SFS for the SV and on its frequency.

### 2.6 Fay and Wu’s *H*

Fay and Wu’s *H* (Fay and Wu (2000)) is the difference of an estimator 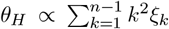, which emphasizes high-frequency derived alleles, and Tajima’s *π*. In regions covered by SVs, 𝔼_*SV*_ [*H*|*i*]≠ 0 even under neutrality.

## 3 Results

### 3.1 Exact conditional SFS under complete linkage

Under neutrality, the expected number of mutations in each frequency class is proportional to the expected total branch length subtending *k* leaves. However, the structure of the spectrum is complicated by the relation between these leaves and the ones containing the derived SV allele.

In this paper, we assume a biallelic SV with a single origin of the derived sequence. In this case, under complete linkage, five subspectra can be recognised, depending on the relation between the mutation and the SV (Ferretti et al. (2018)):

- strictly nested (*sn*): the mutation is present in a proper subset of sequences containing the derived SV allele;
- co-occurring (*co*): the mutation is present in all sequences containing the derived SV allele;
- enclosing (*en*): the derived SV allele is present in a propersubset of sequences containing the mutation;
- complementary (*cm*): each sequence contains either the mutation or derived SV allele;
- strictly disjoint (*sd*): the mutation is present in a proper subset of sequences that do not contain the derived SV allele.

### 3.2 General form of spectrum for linked flanking regions

Assuming that both alleles of the SV can be properly detected and assigned to a sequence, the spectrum for linked flanking regions contains all components, i.e.

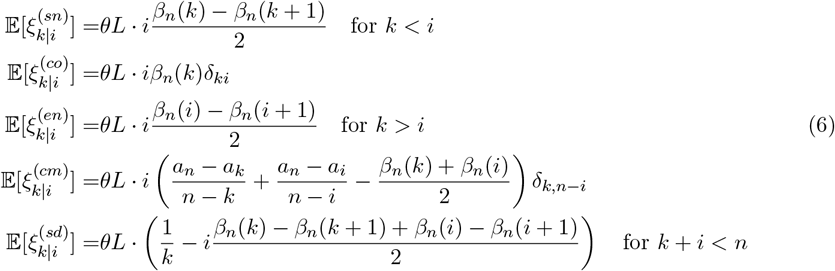

where 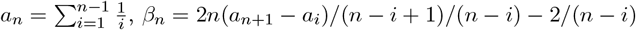, and *θ* is the standard neutral mutation rate parameter per base and *L* is the length of the sequence (Ferretti *et al*. (2018)).

The full spectrum is therefore

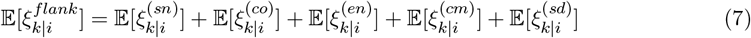

irrespective of the type of SV, provided that it is biallelic and single-origin.

### 3.3 Inversions

For inversions, we assume that sites are present on both arrangements and are perfectly associated with arrangement status. Mutations arising on either background are observable in the aligned sequence. The conditional SFS contains all the components

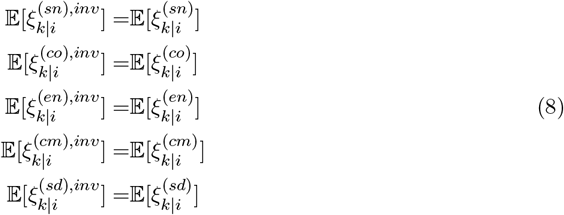

and results in the same overall spectrum as their flanking regions (Giner-Delgado *et al*. (2019)):

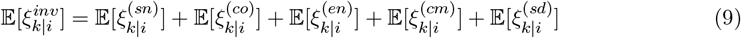

illustrated in Figure 1.

**Figure 1.**
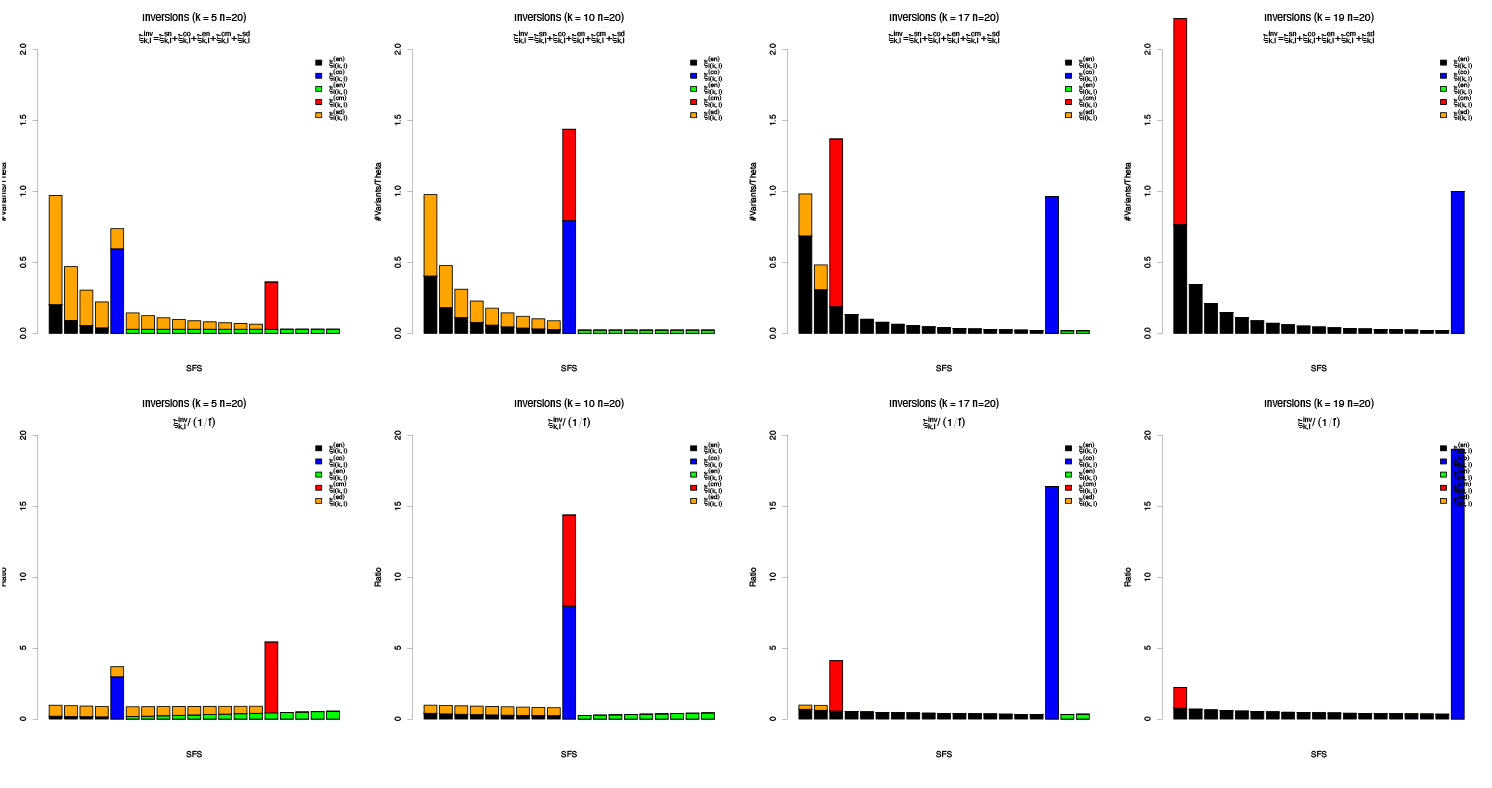
SFS associated to a inversion at different frequencies. Sample spectrum 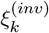 (first row) and Nawa-Tajima statistics, i.e. ratio of sample spectrum compared to the neutral baseline *ξ*_*k*_ (second row).

### 3.4 Deletions

For deletions, the sequence within the deleted segment is present only on the ancestral SV background. For analyses restricted to that segment, observed SNPs arise only on lineages carrying the ancestral SV allele.

The spectrum effectively contains just a single component, the spectrum of frequencies *k* ∈ [1, *n* − *i* − 1] among the *n* − *i* surviving sequences:

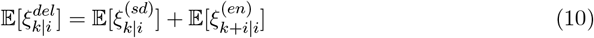

as illustrated in Figure 2.

**Figure 2.**
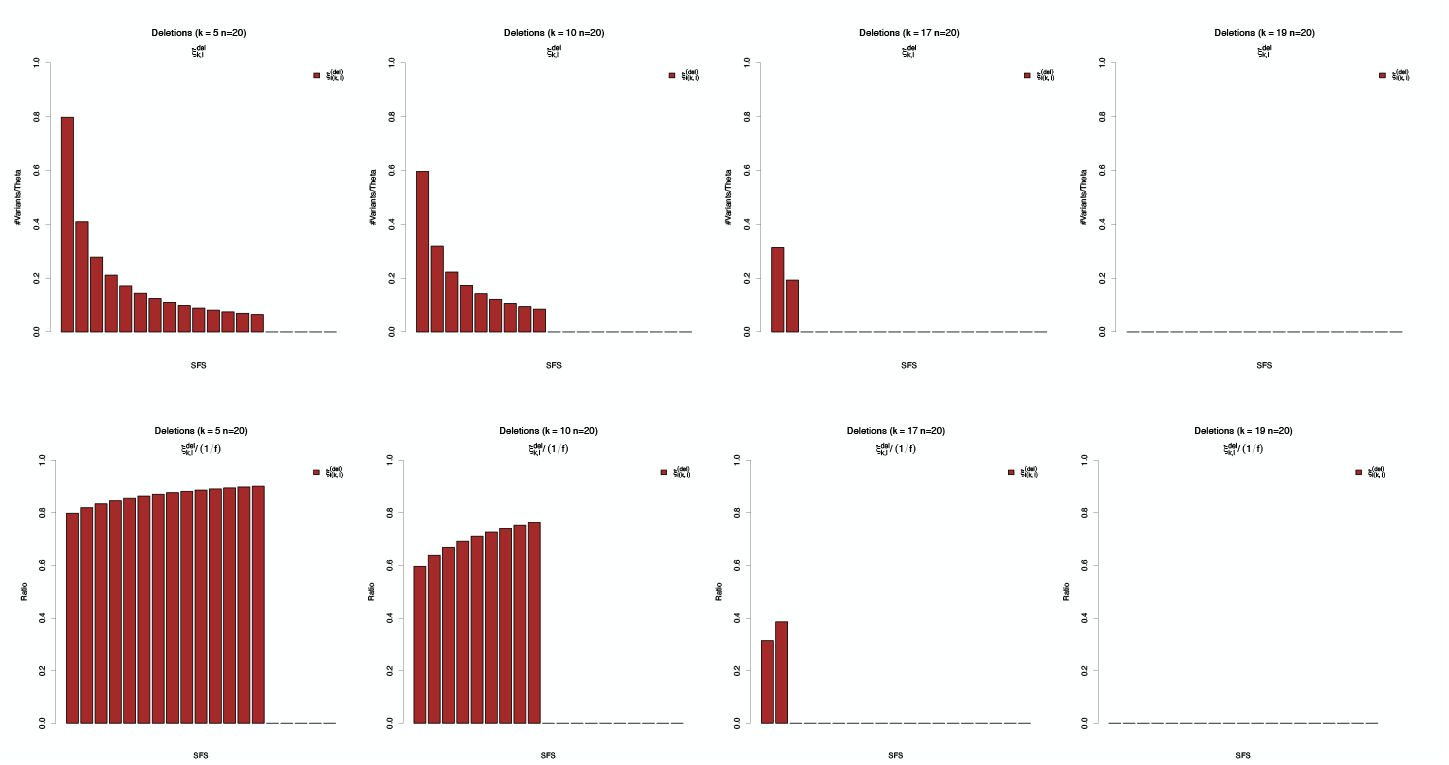
SFS associated to a deletion at different frequencies. Sample spectrum 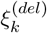 (first row) and Nawa-Tajima statistics, i.e. ratio of sample spectrum compared to the neutral baseline *ξ*_*k*_ (second row).

### 3.5 Insertions

For insertions, insertion-unique sequence is present only on the derived SV background. For analyses restricted to insertion-unique sequence, within the insertion there is just a single component, the spectrum of frequencies *k* ∈ [1, *i* − 1] among the *i* inserted sequences:

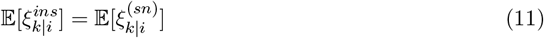

illustrated in Figure 3.

**Figure 3.**
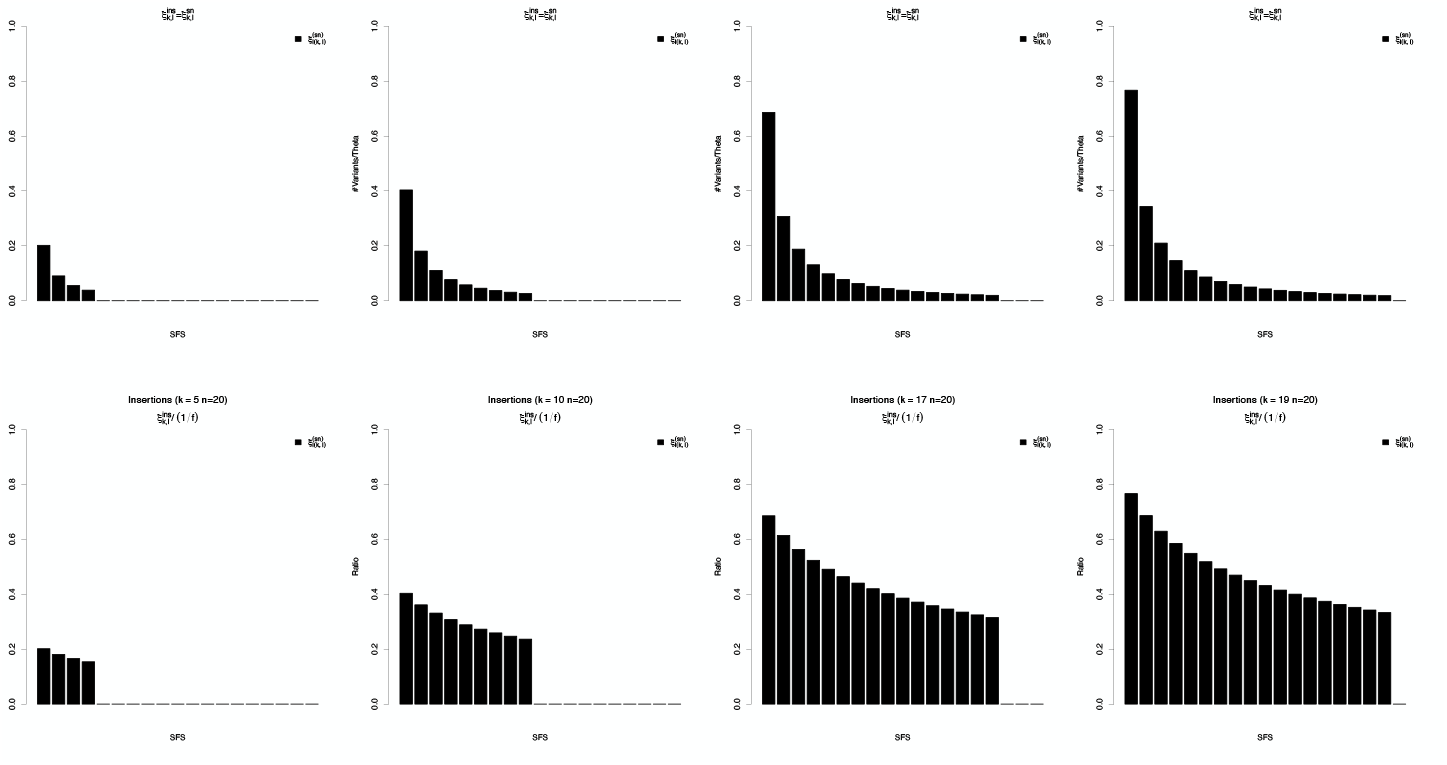
SFS associated to a insertion at different frequencies. Sample spectrum 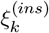 (first row) and Nawa-Tajima statistics, i.e. ratio of sample spectrum compared to the neutral baseline *ξ*_*k*_ (second row).

Note that if the origin of the inserted sequence is unknown, it is usually impossible to polarise the nucleotide polymorphisms and the only possible definition for the spectrum is the folded one:

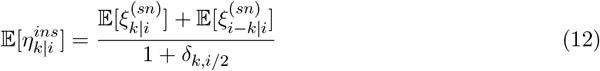

### 3.6 Introgressions

For introgressions, we model the SV allele as being associated with ancestry from a diverged source population, resulting in two clearly divergent backgrounds and deeper genealogical branches on the SV background. The introgression-conditional spectrum generally depends on an additional parameter, i.e. the divergence per base of each allele from their common ancestor. We denote by *D*_*i*_ the apparent divergence between the common ancestor and the introgressed allele, and by *D*_*a*_ the apparent divergence between the common ancestor and the original allele.

Critically, it is usually impossible to unambiguously assign any fixed difference between ancestral and introgressed alleles to be either a result of ancestral divergence or a recent polymorphism fixed in one of the two backgrounds. This means that on one hand, the spectrum contains only two meaningful components for population genetic analyses, the internal and external components with respect to the introgression. The spectra of these components correspond to the spectra of an insertion and a deletion respectively:

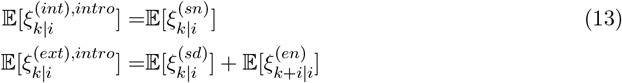

On the other hand, to assess the overall impact of the introgression on diversity and neutrality tests, we can define the full spectrum of an introgression as the apparent distribution in frequency of all detectable nucleotide polymorphisms within the introgressed region:

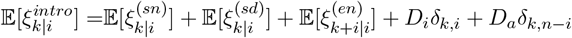

which includes the (polarised) fixed differences between introgressed and ancestral allele.

If the origin of the introgressed sequence is unknown, it is usually impossible to polarise the nucleotide polymorphisms and the spectrum is folded as for insertions.

### 3.7 SV-induced bias in estimators of diversity

We show the impact of SVs on Watterson’s *θ*_*W*_ and Tajima’s *π* estimators of genetic diversity in the region of the SV in Figures 4. The plots show how the impact of the presence of the SV depends both on its type and frequency.

**Figure 4.**
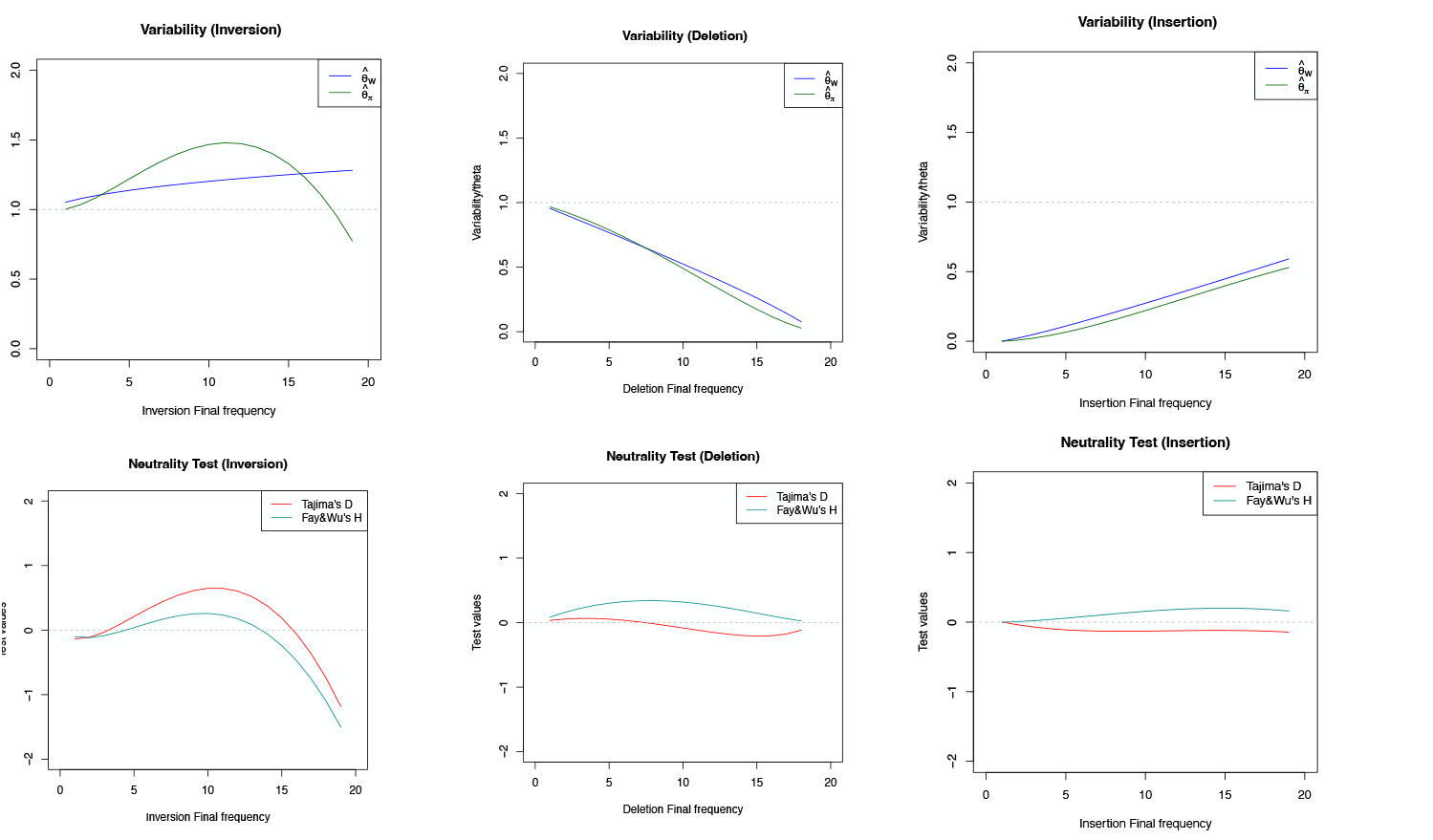
Estimators of Variability and neutrality for inversions, deletions and insertions. first row: *θ*_*W*_ and *π* estimators; second row: and neutrality tests (Tajima’s *D* and Fay and Wu’s *H*).

For introgressions and inversions, the presence of intermediate- or high-frequency variants greatly increases the estimated genetic variability compared to the standard baseline. Watterson’s *θ*_*W*_ is especially prone to be biased upward because of divergence between original and introgressed backgrounds, as expected. Estimated variability only decreases for very high frequency inversions if estimated from pairwise nucleotide diversity *π*.

Both insertions and deletions result in a decrease in estimated variability compared to the standard baseline, even if the estimators already account for the reduced number of sequences. This reduction in estimated variability is especially pronounced for high-frequency deletions and low-frequency insertions.

## 4 SV-induced bias in neutrality tests

In Figure 4 we show the impact of SVs on classical neutrality tests, such as Tajima’s *D* and Fay and Wu’s *H*, when computed in the region covered by the SV. These tests are defined as centered, i.e. their null result is close to 0. The plots show how the deviations from 0 due the presence of the SV depend both on type and frequency of the SV.

Introgressions and inversions of intermediate frequency are generally characterised by an excess of intermediate-frequency mutations, and therefore a positive bias in both *D* and *H* tests. On the other hand, when these SVs reach high frequency, they lead to an excess of rare ancestral alleles, pushing both Tajima’s *D* and Fay and Wu’s *H* towards negative values.

The spectrum of insertions and deletions can be understood as a results of a recent population expansion (for insertions) or contraction (for deletions). Intuitively, this should result in an excess of low-frequency alleles within inserted sequences, leading to *D* < 0 and *H* > 0 for insertions. Similarly, we expect a depletion of low-frequency alleles within deletions, leading to *D* > 0 and *H* < 0 for regions covered by deletions. The analytical results confirm these expectations for the shape of the spectrum, but the expected values of the tests are actually scarcely affected, i.e. Tajima’s *D* and Fay and Wu’s *H* are approximately centered for indels.

### 4.1 SV-aware adjustment of estimators and tests

The conditional expectations derived above provide a natural route to SV-aware adjustments. However, there are multiple possible avenues for these adjustments, each leading to slightly different tests and estimators.

For example, any linear estimator of genetic diversity can be written in the form (Achaz (2009); Ferretti et al. (2010))

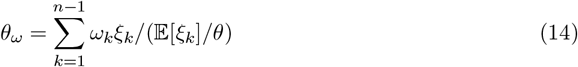

with 𝔼 [*ξ*_*k*_] = *θL/k*. There are at least two options to adjust this estimator to be applied to the region covered by a SV of known frequency *i*. The first one is to redefine it using the appropriate null spectrum that accounts for the SV:

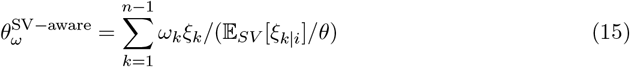

and the second one is to renormalise the estimator, rescale the whole expression by a constant:

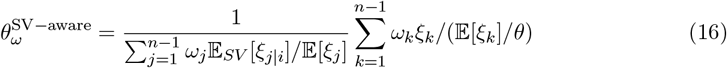

Similarly, neutrality tests are based on differences between two unbiased estimators, and can be appropriately centered by choosing any of the above corrections for the two estimators. The first choice can be interpreted as being more focused on the pattern of relative deviations from the expected spectrum, and the second one more focused on absolute deviations.

The actual difference between different alternative unbiased/centered versions of these estimators and tests is unlikely to be relevant for practical applications. We leave the detailed study of optimal correction, calibration, and power under specific alternatives as an avenue for future work.

## 5 Discussion

Under complete linkage, polymorphic SVs induce frequency-dependent distortions of the neutral SFS that depend on the nature of the SV. These distortions propagate directly to diversity estimators and neutrality tests, producing systematic bias under neutrality. Treating inversions, deletions, insertions, and introgressions separately clarifies how the sources of bias differ: inversions primarily affect the genealogical structure, mutations in deletions and insertions are present on ancestral and derived background-specific site presence and therefore have a very different expected spectrum that accounts for the growth/decrease in frequency of the background, and introgressions add deeper ancestry components parameterized by divergence and admixture histories.

Our analysis makes several simplifying assumptions that delimit its scope. First, we assume complete linkage between the SV and analyzed sites. This is appropriate for tightly linked regions and provides exact theoretical results, but real genomes exhibit some recombination or gene conversion that will partially decouple backgrounds. Our results could be extended by incorporating partial linkage (recombination or gene conversion) and characterizing how bias decays with distance from the SV. Second, we consider simple SV architectures. We treat SVs as biallelic and do not model multi-allelic variants, nested SVs, tandem repeats with complex mutation processes, or SV clusters with correlated breakpoints. It is possible to extend our framework to obtain approximate results for more complex multi-allelic SVs; however, including multi-copy insertions and recurrent SV mutation presents significant challenges (Giner-Delgado *et al*. (2019)). Third, we assume that deletions/insertions are properly called and genotyped. In empirical settings, mapping, alignment, and genotype calling biases can further distort observed spectra in ways not captured here (Ahsan et al. (2023)). Fourth, the introgressions are modelled in a simplified way that introduces additional parameters (i.e. divergence times). A more complete analysis would integrate over more complex plausible histories, including potential admixture, at the price of adding extra parameters. Fifth, we consider a standard demographic baseline. These results could be extended to more general demographic histories with variable population size (Griffiths and Tavaré (1998)) at the price of a significant increase in the complexity of the formalism (Živkovic and Wiehe (2008)).

Finally, our results could be extended to the case of selection acting on the SV or on linked sites, which will further shape the conditional SFS. An exact treatment would be a welcome advancement but is likely to require additional theory, including joint modeling of SV frequency dynamics and linked neutral variation, enabling inference on SV age and selection from conditional SFS patterns.

